# Sensitive Detection of Protein Binding to the Plasma Membrane with Dual-Color Z-Scan Fluorescence

**DOI:** 10.1101/766436

**Authors:** I. Angert, S.R. Karuka, J. Hennen, Y. Chen, J.P. Albanesi, L.M. Mansky, J.D. Mueller

## Abstract

Delicate and transitory protein engagement at the plasma membrane (PM) is crucial to a broad range of cellular functions including cell motility, signal transduction, and virus replication. Here we describe a dual color (DC) extension of the fluorescence z-scan technique which has proven successful for quantification of peripheral membrane protein binding to the PM in living cells. We demonstrate that the co-expression of a second distinctly colored fluorescent protein provides a soluble reference species, which delineates the extent of the cell cytoplasm and lowers the detection threshold of z-scan PM binding measurements by an order of magnitude. DC z-scan generates an intensity profile for each detection channel that contains information on the axial distribution of the peripheral membrane and reference protein. Fit models for DC z-scan are developed and verified using simple model systems. Next, we apply the quantitative DC z-scan technique to investigate the binding of two peripheral membrane protein systems for which previous z-scan studies failed to detect binding: human immunodeficiency virus type 1 (HIV-1) matrix (MA) protein and lipidation-deficient mutants of the fibroblast growth factor receptor substrate 2α. Our findings show that these mutations severely disrupt PM association of fibroblast growth factor receptor substrate 2α but do not eliminate it. We further detected binding of HIV-1 MA to the PM using DC z-scan. Interestingly, our data indicate that HIV-1 MA binds cooperatively to the PM with a dissociation coefficient of *K*_*d*_ ~16 μM and Hill coefficient of *n* ~2.

**SIGNIFICANCE:** Protein binding to the plasma membrane of cells plays an important role in a multitude of cell functions and disease processes. Quantitative binding studies of protein/membrane interactions are almost exclusively limited to in vitro systems and may produce results that poorly mimic the authentic interactions in living cells. We report quantitative measurements of plasma membrane binding directly in living cells by using dual color z-scan fluorescence, which improves the detection threshold by an order of magnitude compared to our previous single color technique. This advance allowed us to examine the role of mutations on binding affinity and identify the presence of cooperative binding in protein systems with relevance to HIV/AIDS and cancer biology.

## INTRODUCTION

The association of peripheral membrane proteins with the plasma membrane (PM) is a crucial and vital step in numerous cellular pathways, such as cell differentiation (1), lipid metabolism (2), signal transduction (3), and oncogenesis (4). Proteins may be recruited to the PM through interactions with integral membrane proteins or by direct interaction with membrane lipids (5, 6). Because the ability to bind rapidly and reversibly to membranes is essential for the function of peripheral membrane proteins, their membrane affinity is a biophysical property of fundamental interest. Membrane binding studies are routinely performed using in vitro assays utilizing liposomes, supported lipid bilayers, or lipid monolayers (7–9). However, extrapolating the results of such studies to proteins in living cells is challenging because the in vitro environment cannot faithfully reproduce the native conditions that give rise to the complex organization and dynamic behavior of cellular membranes (10). Therefore, direct measurements of membrane binding in living cells are needed to explore the binding process in its natural environment and critically examine observations derived from in vitro work.

We recently succeeded in directly measuring the PM binding curve of EGFP-tagged peripheral membrane proteins in living cells by single color (SC) z-scan (11). An axial scan of the focal spot generated by two-photon excitation through a chosen cytoplasmic location produces a z-scan intensity profile. This profile is the convolution of the two-photon point spread function (PSF) of the instrument with the axial concentration profile of the labeled protein along the scan trajectory (12). A population of PM-bound protein leads to the appearance of intensity peaks along the scan path at the location of the bottom and top PM. Modeling of the intensity profile separates cytoplasmic and membrane-bound fluorescence components, which allows for the determination of the protein concentration in the cytoplasm and at the membrane (13). We successfully applied this approach to characterize PM binding of human T-cell leukemia virus type 1 (HTLV-1) matrix (MA) protein in live cells, and identified the binding affinity as well as the saturating binding concentration of the protein (11). However, this method failed to detect PM binding of the matrix protein of a different retrovirus, human immunodeficiency virus type-1 (HIV-1) (14). Because HIV-1 MA interacts much more weakly with the PM than HTLV-1 MA, the membrane-bound population of HIV-1 MA is below the detection threshold of the SC z-scan method. In general, detection of PM binding by SC z-scan is limited to proteins with relatively high binding affinity or high binding site density at the PM.

To overcome this limitation of SC z-scan we co-expressed the EGFP-labeled membrane-binding protein together with mCherry to provide a reference marker of the cytoplasmic space. The z-scan intensity profile of mCherry furnishes independent information on the starting and ending point of the cytoplasm, thereby improving the resolution of small fluorescence intensity contributions from the PM-bound EGFP-labeled protein. In this study we demonstrate that the simultaneous collection of the intensity profiles of the EGFP-labeled protein and mCherry in two separate detection channels increases the ability to resolve PM binding by an order of magnitude. We establish a framework for analyzing DC z-scan measurements and verify it experimentally. Control experiments were performed to explore the sensitivity threshold of detecting a PM-bound protein population from DC z-scan profiles.

Here we applied DC z-scan to characterize PM binding of the fibroblast growth factor receptor substrate 2α (FRS2α). It was recently established that FRS2α not only is myristoylated near its N terminus but is also doubly palmitoylated (15). Removing the palmitoyl groups weakened PM binding to the detection limit of the previously described SC z-scan technique (15). By applying DC z-scan we were able to identify residual PM binding in the absence of palmitoylation and compare this binding to that of wildtype (WT) FRS2α. DC z-scan was further applied to characterize PM-binding of HIV-1 MA in live cells. Interestingly, a simple Langmuir-isotherm model was inconsistent with the experimental data. Our results indicate that HIV-1 MA binds cooperatively to the PM.

## MATERIALS AND METHODS

### Sample Preparation, Cell Culture & Expression Vectors

U-2 OS and HeLa cell lines were cultured in DMEM supplemented with 10% FBS (Hycolone Laboratories, Logan, UT). For measurement, cells were plated into chambered 8-well slides (Cellvis, Mountain View, CA) and transfected with the indicated plasmids using GenJet (SignaGen Laboratories, Rockville, MD) transfection reagent according to the manufacturer instructions. Transfections were performed 12-24hr before measurement with cells at ~50% confluency. Cell media was exchanged for phosphate buffered saline immediately before measurement. Chambered slides were held at room temperature on the microscope during measurement. In vitro experiments utilized EGFP and mCherry purified from *E. coli* as previously described (16).

HIV-1 MA was amplified from HIV-1 Gag (17) with a 5’ primer that encoded a XhoI restriction site and 3’ primer that encoded a EcoRI site. The PCR product was digested and ligated into pEGFP-N1 (clonetech, CA) to generate the HIV-1 MA EGFP vector. HIV-1 MA^G2A^-EGFP was made using the QuikChange XL Site-Directed Mutagenesis Kit (Stratagene, La Jolla, CA). HIV-1 MA^G2A-ΔHBR^-EGFP was generated from HIV-1 MA^G2A^-EGFP by deleting amino acids 18-32 using the Q5 Site-Directed Mutagenesis Kit (New England Biolabs, Ipswich, MA). pEGFP-N1 and pEYFP-N1 were obtained from Clonetech (Mountainview, CA). All other plasmids have been previously described: mCherry and EGFP-mCherry (18); HTLV-1 MA-EGFP (14); FRS2α-EGFP, FRS2α^G2A^-EGFP and FRS2α^C4,5S^-EGFP (15).

### Supported Lipid Bilayer

Supported lipid bilayers consisted of 1-palmitoyl-2-oleoyl-glycero-3-phosphocholine (POPC; Avanti Polar Lipids, Alabaster, AL) with a small admixture of Texas Red and BODIPY FL conjugated to DHPE lipids (ThermoFisher Scientific, Waltham, MA) to provide fluorescent labels. The lipids were mixed in chloroform at 1:3300 ratio (BODIPY FL DHPE : POPC) and 1:1100 ratio (Texas Red DHPE : POPC) and dried under nitrogen flow. Dried lipids were mixed with PBS to form large unilamellar vesicles, which were then extruded to generate small unilamellar vesicles (SUVs) using a 100 nm filter and the NanoSizer MINI Liposome Extruder (T & T Scientific, Knoxville, TN) according to manufacturer instructions. Coverslips were cleaned by brief exposure to a butane flame followed by ~1 hr etch in 50/50 w/w KOH solution. The SUV solution was added to each coverslip for 15 min, followed by a wash with PBS to remove any free floating SUVs.

### Instrumentation and Measurement

DC z-scan measurements were performed on a Zeiss Axiovert 200 microscope modified for two-photon excitation. 1000nm excitation light provided by a mode-locked Ti:S laser (MaiTai or Tsunami models, Spectra Physics, Santa Clara, CA) is focused to a diffraction limited spot at the sample by a 1.2 numerical aperture water immersion objective (C-Apochromat, Zeiss, Oberkochen, Germany). Fluorescence emission light is collected by the same objective and separated from excitation light by a short-pass dichroic (740DCSPXR, Chroma Technology, Bellows Falls, VT) before being split into red and green detection channels by a 580nm long-pass dichroic filter (FF580-FDi01; Semrock, Rochester, NY) and detected by single photon counting detectors (hybrid PMT; HPM-100-40, Becker and Hickl, Berlin, Germany). A 515/50nm band pass filter is placed immediately before the green channel detector to eliminate Fresnel-reflected mCherry light from the green channel.

To perform DC z-scan measurements, samples were mounted on a stage with piezo-driven z-axis (PZ2000, ASI, Eugene OR) that is controlled by a voltage signal originating from an arbitrary waveform generator (model 33250A, Agilent Technologies, Santa Clara, CA). DC z-scans were performed by driving the piezo stage with a triangle waveform with a period *T* and peak-to-peak displacement *Z*_*pp*_. All DC z-scan measurements were performed with *T* = 10 s and *Z*_*pp*_ = 24 μm. Thus, each 10 s period yields two z-scan traces, each 5 s or 24 μm in length; the first trace corresponds to motion of the PSF upward through the sample and the second to motion downward through the sample.

### Modeling of z-scan traces

The PSF of our instrument is well-approximated by the modified squared Gaussian-Lorentzian (mGL) PSF with parameters typically in the range of z_*R*_ = 0.70 ± 0.15 μm, *y* = 1.3 ± 0.2, and *w*_0_ =0.47 ± 0.05 μm. The precise *z*_*R*_ and *y* parameters are determined by a daily calibration with an uncertainty of 1 – 5% as previously described (12).

This results in a typical PSF volume of *V*_∞_ = 0.3 ± 0.1 μm^3^ and a cross-sectional area of *A*_0_ = 0.17± 0.04 μm^2^. The radially integrated PSF, 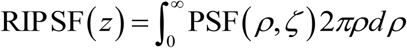 provides the foundation for modeling z-scan intensity profiles (12). The normalized observation volume is defined by definite integration of the RIPSF function (11),

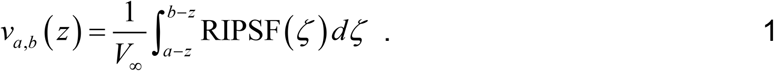

For the special case of a δ-layer Eq. 1 reduces to 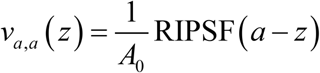 (11).

Analytical formulas (11) of the volume function are listed in the Supporting Material.

### Characterization of z-scan by simulation

DC z-scan traces were modeled based on a δ-s-δ layer for the EGFP labeled protein and an s layer for mCherry (Eq. 13) with added Poisson noise to account for the photon counting process. The fluorescence intensity amplitudes of the membrane and cytosolic protein populations were chosen as described in the Results section. The simulated DC z-scan data were fit using the same procedure as applied to experimental data. Separate simulations to characterize the SC z-scan method were performed similarly except only green channel intensity traces were simulated and fit. All summary statistics presented in the text represent the result of fits to 1000 realizations of each z-scan trace. Fits that were observed to become stuck in local minima were identified based on elevated reduced chi-square and removed from the data set before the calculation of summary statistics. All modeled traces were calculated assuming an mGL PSF with parameters *z*_*R*_ = 0.75 μm and *y* = 1.35.

### Membrane binding equilibria

To construct binding equilibria, the cytoplasmic (*f*_*C*_) and membrane-bound (*f*_*M*_) fluorescence amplitudes (Eq. 8) were converted into cytoplasmic concentrations *c*_*C*_ and membrane-bound surface density *σ*_*M*_ (13),

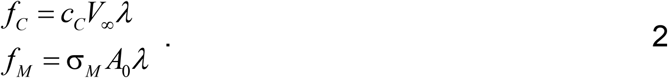

The brightness *λ* of the label EGFP was determined by calibration experiments as previously reported (12).

Membrane binding equilibria can be modeled by the Hill equation, which depends on three parameters, the dissociation coefficient *K*_*d*_, the saturation density *σ*_0_, and the Hill exponent *n*,

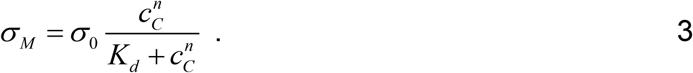

The membrane intensity fraction (Eq. 14) of the Hill equation is identified by

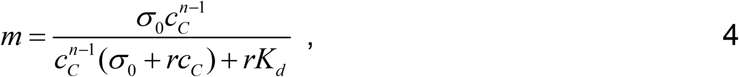

where *r* is the ratio of the PSF volume to its cross-sectional area,

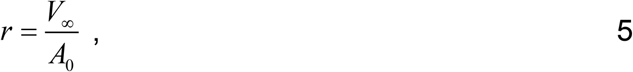

which for our instrument is given by 1.6 μm. The Hill equation reduces to the Langmuir isotherm for the case *n* =1, which describes non-cooperative membrane binding. Experimental data were first fit to a Langmuir isotherm and subsequently to the Hill equation in cases where the Langmuir isotherm failed to adequately describe the data. For a Langmuir isotherm the maximum value of *m*, which we denote as *m*_0_, is obtained as the cytoplasmic concentration approaches zero (11). The ratio of the binding parameters *K*_*d*_ and *σ*_0_ are related to *m*_0_ by

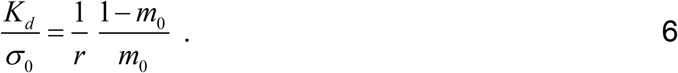

## RESULTS

The building block of SC z-scan analysis is the slab layer (s layer). This horizontal layer extends from height *a* to *b* with uniform protein concentration *c*_0_ within the slab and zero outside the layer (Fig. 1A),

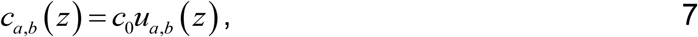

with the boxcar function *u*_*a,b*_(*z*), which is equal to 1 for *a* ≤ *z* ≤ *b* and 0 otherwise. Two limiting cases of the s layer are of interest. First, the infinitesimally thin layer or delta (δ) layer located at *a* (Fig. 1A) represents the limit *b* → *a* with concentration profile *c*_*a*,*a*_(*z*) = *σ*_0_*u*_*a*,*a*_(*z*) = *σ*_0_ δ(*z* − *a*), where *σ*_0_ is the surface concentration (13) (Fig. 1B). Second, the semi-infinite layer represents the case where one of the boundaries goes to infinity, i.e. *c*_*a,∞*_(*z*) (Fig. 1C).

**Figure 1.**
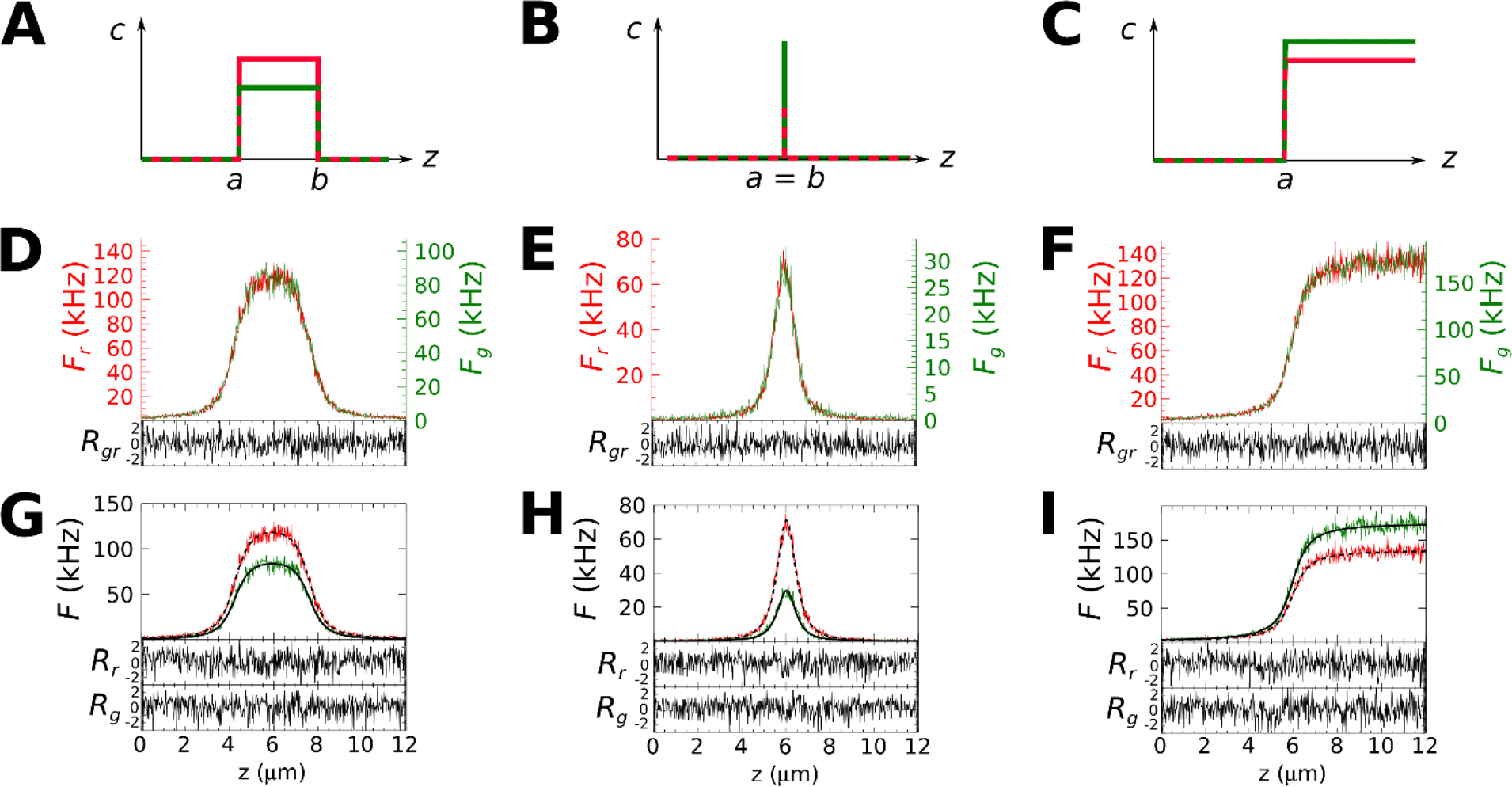
DC z-scan of three sample layers. The two detection channels of the DC z-scan are represented by the green and the red color. An illustration of the fluorescence profile as a function of axial height *z* are shown for a slab layer (A), a delta layer (B), and the semi-infinite layer (C). The start and end height of each layer is marked by *a* and *b*, respectively. The experimental intensity profiles *F*_*g*_(*z*) of the green channel are scaled to the intensity profile *F*_*r*_(*z*) of the red channel and shown for the slab (D), delta (E), and semi-infinite layer (F) together with the normalized residuals, *R*_*gr*_ of the scaling. Fits of the unscaled intensity profiles of the slab (G), delta (H), and semi-infinite layer (I) to the corresponding DC z-scan model are shown with normalized residuals, *R*_*g*_ and *R*_*r*_ corresponding to the green and red channel fits, respectively.

An axial scan of the two-photon PSF through a concentration layer *c*_*a*,*b*_(*z*) gives rise to a fluorescence intensity profile *F*(*z*), which reflects the spatial overlap of the PSF with the concentration profile. We previously showed that the z-scan intensity profile is modeled by (12)

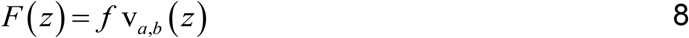

with *f* representing the amplitude, which is proportional to the concentration *c*_0_ of the layer, and the volume function v_*a*,*b*_(*z*), which is derived from the convolution of the PSF with the sample layer (12). Formulas describing v_*a*,*b*_(*z*) are listed in the Supplemental information. Straightforward extension of the SC z-scan model to two colors requires that the volume function v_*a*,*b*_(*z*) remain identical for each detection channel.

To experimentally test this proposition, we performed DC z-scans on model systems. First, we selected a cytoplasmic location of U-2 OS cells co-expressing EGFP and mCherry (CH) to mimic a slab layer. A DC z-scan measurement determined the intensity profiles of the green and red channel, *F*_*g*_(*z*) and *F*_*r*_(*z*). To simplify visual comparison of both curves we scaled *F_g_*(*z*) by a factor α to match the amplitude of the red intensity curve, 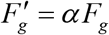 (Fig. 1D). The normalized residuals between the green and red curve are flat with a reduced chi-square 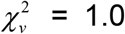 (Fig. 1D, lower panel). The absence of a mismatch between both intensity curves indicates that the volume function v_*a*,*b*_(*z*) is the same for each channel within experimental uncertainty.

Next, we performed DC z-scans on a delta layer, which was realized by a supported lipid bilayer containing Texas Red and BODIPY FL conjugated lipids. We again scaled *F*_*g*_(*z*) to match the amplitude of *F*_*r*_(*z*) (Fig. 1E) and calculated the normalized residuals (Fig. 1E, bottom panel). Again, the scaled intensity traces are indistinguishable 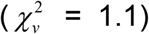. Finally, a mixture of purified EGFP and mCherry was added to a well-slide to create a semi-infinite layer. The scaled intensity profiles of both channels acquired with DC z-scan coincide (Fig. 1F) with flat residuals (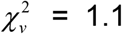, Fig. 1F, lower panel).

These results suggest that modeling of DC z-scan intensity profiles is a straightforward extension of Eq. 8,

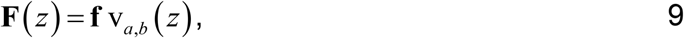

where **F**(*z*)=(*F*_*g*_(*z*), *F*_*r*_(*z*)) and **f** =(*f*_*g*_, *f*_*r*_) represent the vector notation of the intensity profile and amplitude of the green and red detection channel. The volume function v_*a*,*b*_(*z*) depends on the PSF model. We choose the modified Gaussian-Lorentzian (mGL) PSF, which proved successful in describing single color z-scan intensity profiles (12). To evaluate the DC z-scan model we fit the unscaled intensity profiles of the three model geometries to Eq. 9 using the volume function for the mGL model with fixed PSF parameters. A fit of the intensity profiles of the cytoplasmic layer to a slab model identifies a slab thickness *L* = *b*−*a* of 3.28 μm with good agreement between data and fit (Fig. 1G). Similarly, the intensity profile of the supported lipid bilayer is reproduced by a fit to a delta-layer model (*a* = *b*) (Fig. 1H). Finally, the semi-infinite layer model ( *b*→∞) successfully describes the intensity profiles of the dye mixture solutions (Fig. 1I). Together, these results demonstrate the success of Eq. 9 in modeling the DC z-scan profile of a single layer.

We next consider a single fluorescence species, i.e. EGFP, occupying multiple layers. For single color z-scan the profile from multiple layers is given by the superposition of the intensity profiles of each layer (14). Extending this result to DC z-scan we get

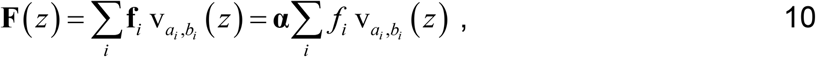

where *a*_*i*_ and *b*_*i*_ are the start and end coordinates of the *i*-th layer with amplitude vector **f**_*i*_. The fluorescence intensity is split into each detection channel as determined by the optical filters and the emission spectrum of the fluorescent protein. This spectral crosstalk ensures that, for a single species, the fluorescence of the green and red channel are proportional to each other. This relation is quantified by the crosstalk vector **α**, which relates the intensity amplitudes of both detection channels, **f**_*i*_ = **α***f*_*i*_, where *f*_*i*_ = *f*_*i,*g**_ + *f*_*i*,*r*_ is the total amplitude of both channels combined. We present a DC z-scan of a cell expressing HTLV-1 MA-EGFP to illustrate multiple layers (Fig. 2). The peripheral membrane protein MA is found both at the bottom and top PM as well as in the cytoplasm as indicated by its concentration profile (Fig. 2A). The two intensity peaks visible in the green channel represent the signal from MA-EGFP bound to the membranes (Fig. 2B). The concentration profile of a peripheral membrane protein is modeled by a δ-s-δ layer model, where the δ layers represent the two membranes and the s layer describes the cytoplasmic space (C) separating the top membrane (TM) and bottom membrane (BM) (14). The DC z-scan profile of the δ-s-δ model for an EGFP-labeled protein is,

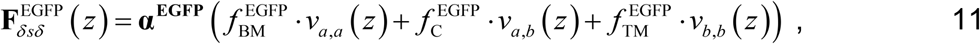

with the membranes located at *z* = *a* and *z* = *b*. The red tail of EGFP’s emission spectrum gives rise to a crosstalk signal in the red detection channel (Fig. 2B) with a crosstalk vector defined by **α**^**EGFP**^ =(1−*α*^EGFP^,*α*^EGFP^). The experimental DC z-scan data are well described by the δ-s-δ model 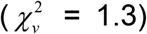 with *α*^EGFP^ = 0.11 (Fig. 2B). The fluorescence contributions from each layer were calculated from the fitted fluorescence intensity amplitudes 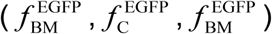 as well as membrane locations (*a*, *b*) and are identified in Fig. 2C.

**Figure 2.**
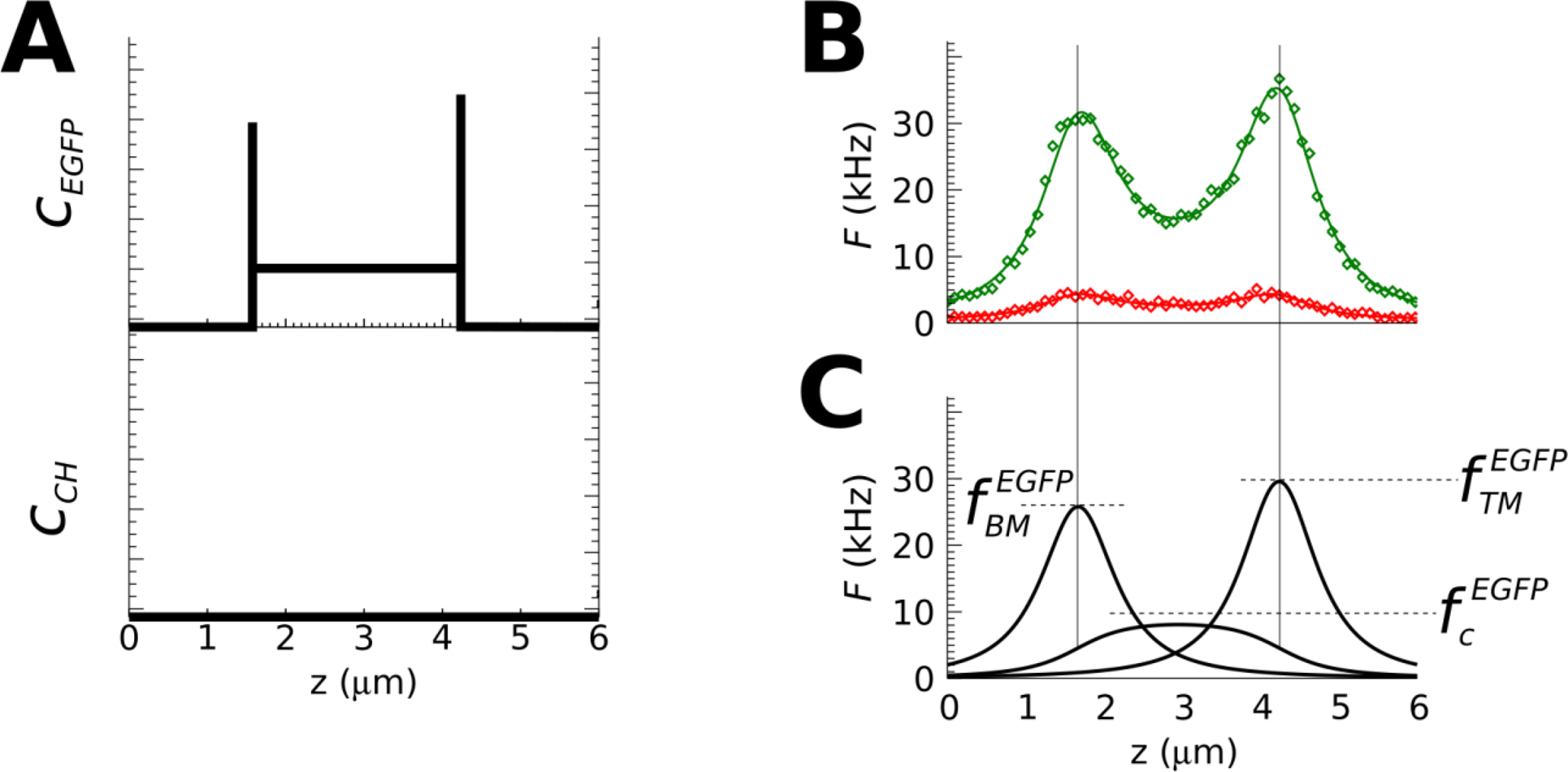
DC z-scan of HTLV-1 MA-EGFP. A) Schematic concentration profile through a cell expressing MA-EGFP without mCherry. The protein is present in the cytoplasm as well as at the upper and lower PM. B) DC intensity profile of the green and red detection channel with fit (solid line) to a δ-s-δ model (Eq.11). C) Decomposition of intensity profile into its cytoplasmic and PM components. The fluorescence amplitudes as well as the location of the membrane layers are determined by the fit.

To improve the accuracy of the fit model we routinely include background above and below the sample. For a sample starting at height *a* and ending at *b* two semi-infinite layers, **f**_↓_ v_−∞,*a*_(*z*) and **f**_↑_ v_*b*,∞_(*z*), are added to the fit model to account for background with DC amplitudes of **f**_↓_ and **f**_↑_. We illustrate the effect of background by considering a δ layer located at *a*, which was experimentally realized by a supported lipid bilayer. Modeling of the DC z-scan profile with a single δ layer, **f**_*δ*_v_*a*,*a*_(*z*), resulted in a poor fit (Fig. S1B). After including background, **f**_↓_v_−∞,*a*_(*z*) + **α**^**EGFP**^*f*_δ_v_*a*,*a*_(*z*) + **f**_↑_v_*a*,∞_(*z*), data and fit are in agreement (Fig. S1A). Although we omit explicit mention of the background component, all data presented in this study were analyzed with background layers added to the fit model where appropriate.

The second color of our study is provided by mCherry. The optical filters are chosen to prevent crosstalk of mCherry emission into the green detection channel, which predicts a crosstalk vector of **α**^**CH**^=(0,1). The lack of crosstalk is confirmed by a cytoplasmic DC z-scan through a cell expressing mCherry (Fig. S2), which was fit by a slab model,

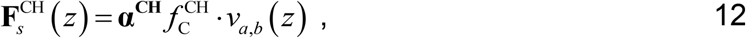

where **α**^**CH**^ is allowed to vary. This fit recovered **α**^**CH**^ = (0.014, 0.986). Because the 1.4% mCherry crosstalk estimated by this fit is generally negligible, we fixed **α**^**CH**^ = (0, 1) in all subsequent DC z-scan fits.

DC z-scan profiles that contain both EGFP and mCherry are described by a superposition of the profiles for EGFP and mCherry, **F**(*z*) = **F**^**EGFP**^(*z*) + **F**^**CH**^(*z*), where each individual profile is modeled by Eq. 10. We are particularly interested in an EGFP-labeled protein that associates with the plasma membrane in the presence of cytoplasmic mCherry. The EGFP z-scan profile follows the δ-s-δ model (Eq. 11), while the mCherry z-scan profile is modeled by the s layer (Eq. 12; Fig.3A). Adding both profiles together yields

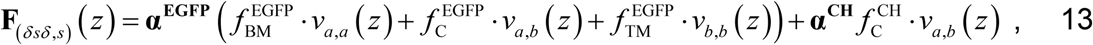

a model described by four amplitudes and the locations *a* and *b* of the membranes. Eq. 13, which we refer to as the δ-s-δ,s model, represents the δ-s-δ model adapted for the present DC z-scan application. This fit model successfully reproduces a DC z-scan profile through the cytoplasm of a U-2 OS cell co-expressing HTLV-1 MA-EGFP and mCherry (Fig. 3B). The decomposition of the green channel signal into its membrane and cytoplasmic component is shown in Fig. 3C. A useful parameter for modeling membrane binding of an EGFP-labeled protein is the membrane intensity fraction (13)

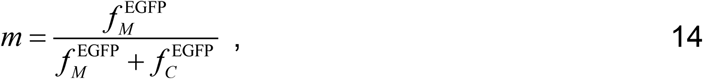

where the subscript *M* either refers to the top (TM) or bottom (BM) membrane. The fluorescence amplitudes at the membrane 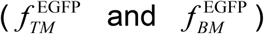 and the cytoplasm 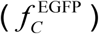 are identified from fitting the DC z-scan intensity profiles to Eq. 13. The fitted DC z-scan of HTLV-1 MA-EGFP (Fig. 3B&C) resulted in *m* of 60% and 56% for the top and bottom membrane, respectively.

We next performed a DC z-scan on a cell expressing the EGFP-labeled matrix protein of HIV-1 in the presence of mCherry (Fig. 3E), which is in stark contrast to the previous DC z-scan. Not only have the peaks in the green-channel profile *F*_*g*_(*z*) disappeared, but there is little difference in the shape of the profiles from both channels (Fig. 3E), indicating the absence of a pronounced membrane-bound HIV-1 MA-EGFP population. Nevertheless, a fit of the DC z-scan traces to Eq. 13 identified a small contribution from a membrane-bound population (Fig. 3F), resulting in membrane intensity fractions of 5.4% and 3.2% for the top and bottom membranes, respectively. Repeated z-scans at the same location and fits to the resulting traces yielded average *m* of 4.1% ± 0.8% and 0.9% ± 0.6% (SEM, *n* = 12) for the top and bottom PMs. These small membrane intensity fractions cannot be recovered by analysis of the green channel z-scan profile *F*_*g*_(*z*) alone. In fact, our previous attempt to detect HIV-1 MA binding to the PM by single color z-scan failed (14), which agrees with earlier results demonstrating that *m* needs to exceed ~10% for single color z-scan analysis to succeed (13).

**Figure 3.**
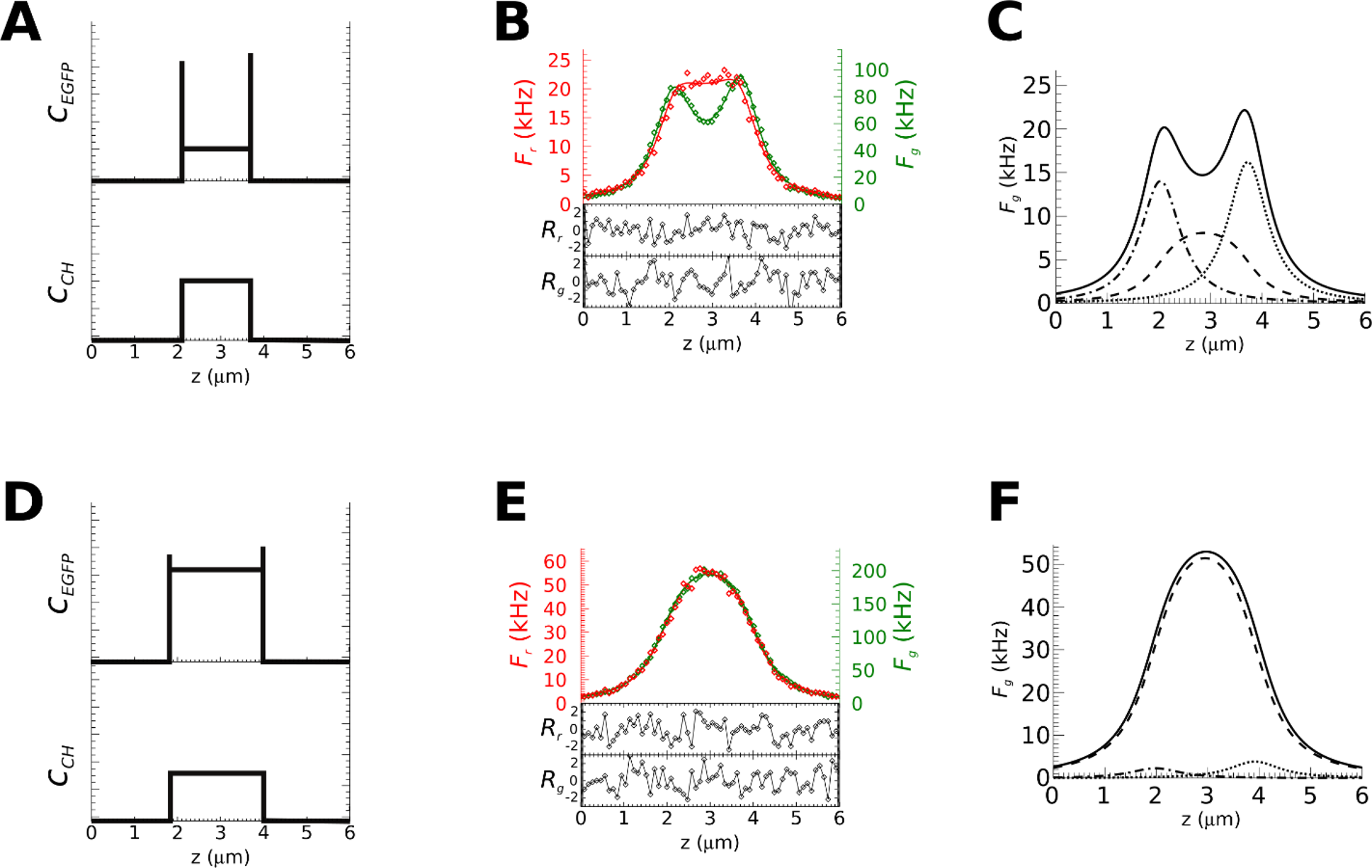
Cytoplasmic DC z-scan of HTLV-1 and of HIV-1 MA-EGFP in the presence of mCherry. A) Schematic concentration profile of HTLV-1 MA-EGFP and mCherry. B) Intensity profile together with fit to a δ-s-δ,s model (Eq. 13). The residuals correspond to 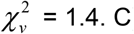 Fit (solid line) of intensity profile for EGFP decomposed into cytoplasmic (dashed line) as well as the upper PM (dotted line) and lower PM contribution (dash-dotted line). D) Schematic concentration profile of HIV-1 MA-EGFP and mCherry. E) Intensity profile together with fit to Eq. 13. The residuals correspond to 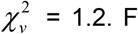 Fit (solid line) of intensity profile for EGFP decomposed into cytoplasmic (dashed line) as well as the upper PM-bound (dotted line) and lower PM-bound contribution (dash-dotted line).

These results suggest that the DC z-scan method has a significantly lower threshold for detecting a membrane-bound population of protein compared to SC z-scan. We turned to modeling to compare the SC and DC z-scan procedure and assess their potential to detect PM-bound fluorescent layers. Z-scan profiles of an EGFP-labeled protein with specific membrane intensity fractions *m*_in_ in the presence or absence of cytoplasmic mCherry were modeled as described in the Methods section. Each calculated z-scan profile included shot noise and was fitted to either the SC δ-s-δ layer model as previously described (14) or the DC δ-s-δ,s layer model (Eq. 13) to recover the fitted *m*_out_ value. The average and standard deviation of *m*_out_ was calculated from fits of *n* = 1000 modeled profiles. To assess whether the DC z-scan procedure recovers correct *m* values, we plotted the average fitted *m*_out_ versus the input value *m*_in_ of the modeled profiles. Because previous SC z-scan studies fitted traces with the constraint that all fluorescent amplitudes be positive, we considered fit procedures with and without this constraint for both SC and DC.

SC z-scan correctly recovered *m*_in_ provided its value was large (*m*_in_ > 10%), but returns biased estimates for small *m*_in_ (Figs. 4A & C). With the constraint of positive fluorescence amplitudes this bias manifested predominantly as a positive offset in recovered *m*_out_ and depends on the overall intensity of the modeled profile (Fig. 4A), while without the constraint the offset could be either positive or negative (Fig. 4C). Similarly, results obtained for modeled DC z-scan profiles with fits constrained to positive amplitudes revealed the presence of bias in the recovered *m*_out_ at small values of *m*_in_ (Fig. 4B). Importantly, once the constraint was removed the DC z-scan fit method recovered unbiased *m*_out_ values within statistical uncertainty (Fig. 4D).

**Figure 4.**
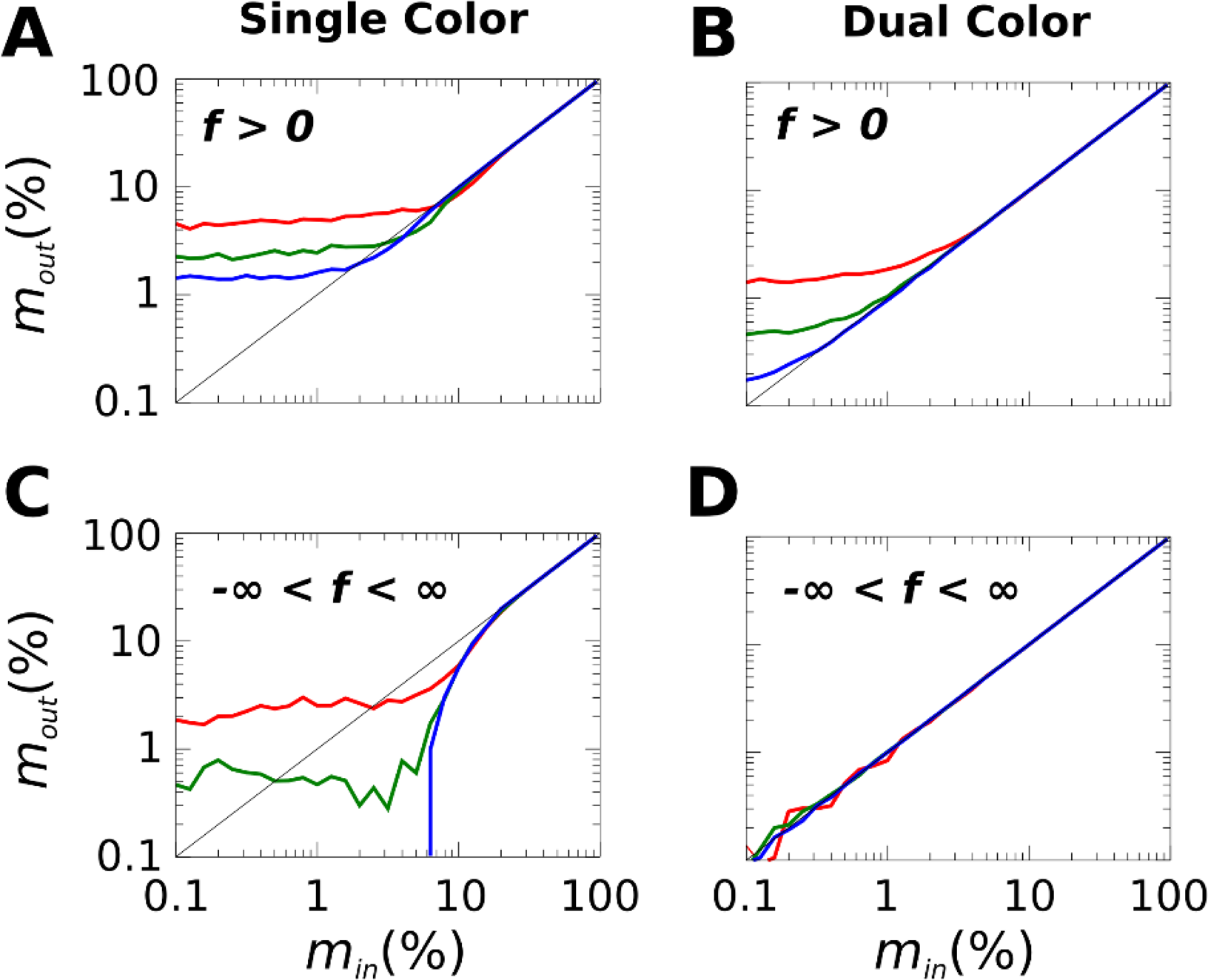
Performance of SC and DC z-scan in recovering the true membrane intensity fraction *m*_in_ from modeling Eq. 13 in the presence of shot noise. The recovered value *m*_out_ from fits of the modeled data to Eq. 13 is plotted vs. the input value *m*_in_ for SC z-scan with fitted fluorescence amplitudes restricted to positive values (Panel A) and unrestricted values (Panel C). The DC z-scan results for restricted (Panel B) and unrestricted (Panel D) fitting of amplitudes are shown for comparison. The black solid line represents *m*_out_ = *m*_in_. The red, green, and blue curve correspond to modeled cytoplasmic intensity amplitudes of 10 kHz, 100 kHz, and 1000 kHz, respectively and a cytoplasm length, *L* = 3μm.

We further compared the standard deviation *s*_*m*_ of the recovered *m*_out_ value for the constrained SC and the unconstrained DC z-scan method as a function of the input value *m*_in_ in Fig. 5A, where each colored line represents a different cytoplasmic intensity. For large membrane intensity fractions *m*_in_ the SC and DC method achieve identical performance, while at low *m*_in_, the DC method outperforms the SC approach by a factor 3 to 6. In general, DC z-scans with higher intensity values lead to lower uncertainty because the relative contribution of the Poisson noise to the signal decreases (13). Quantitatively, we expect *s*_*m*_ to be inversely proportional to the square root of the total number of photons collected, *N*_*γ*_. We are primarily interested in applying DC z-scan to recover small membrane fractions (*m* < 10%) where the contribution of the PM-bound population to the total intensity is small. Under these conditions *s*_*m*_ is approximately independent of *m* (Fig 5A) and the number of photons collected during a z-scan will be proportional to the intensity amplitude and the length of the slab layer, 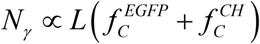. We expect that the statistical uncertainty of *m* is then given by,

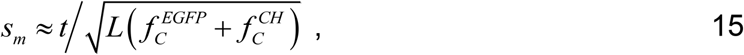

where *t* is a constant that accounts for the shape of the intensity profile. This model provides a means to estimate the expected DC z-scan performance by accounting for *N*_*γ*_. For convenience, we rescaled *s*_*m*_ of the DC z-scan simulations of Fig. 5A to the conditions of the green solid line 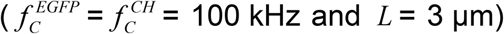, which resulted in overlapping curves (Fig. 5B) and validates this approach. We further repeated the same simulations with a reduced thickness and also with an altered ratio 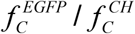 for the three cytoplasmic intensities. The results were rescaled to match the conditions of the green solid line of Fig. 5A to simplify comparison. We note that in all cases the scaled standard uncertainty is close to 1% (Fig. 5B).

**Figure 5.**
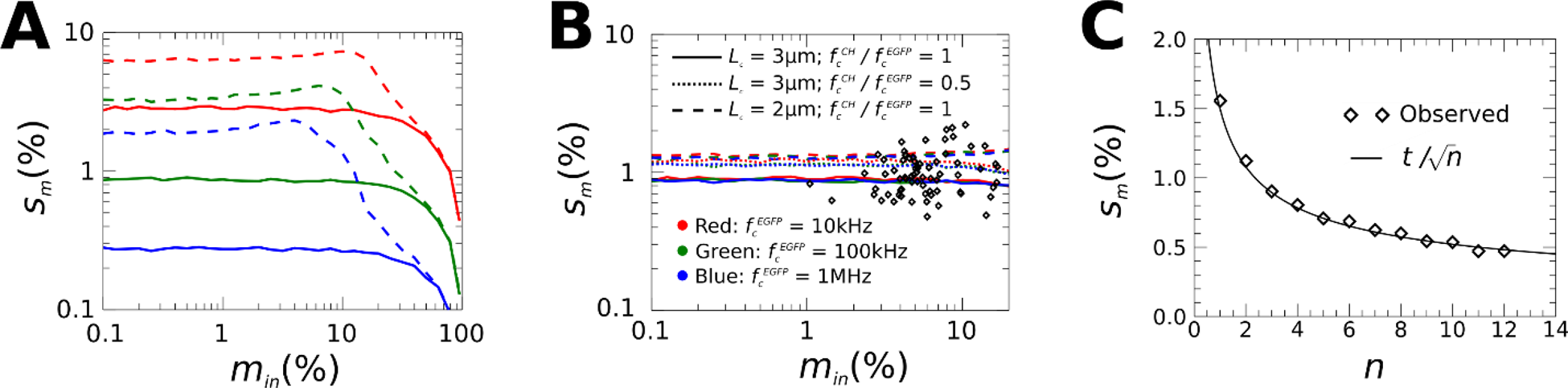
Uncertainty *s*_*m*_ in the recovered membrane intensity fraction *m*. A) The results from simulations of SC z-scan fitted using positive amplitudes is given by the dashed lines. The results from simulations of DC z-scan with unrestricted fitting of amplitudes is represented by the solid lines. The red, green, and blue lines correspond to modeled 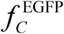 of 10 kHz, 100 kHz, and 1000 kHz, respectively. B) The performance of Eq. 15 is assessed by using this relation to re-scale *s*_*m*_ from simulated data for differing cytoplasmic intensities and lengths so that they match the conditions of the green solid line in A. Experimental results (diamonds) obtained from fitting DC z-scan data of cytoplasmic HIV-1 MA-EGFP followed by re-scaling (using Eq. 15) the observed variance from *n* = 12 repeated scans to match the same reference conditions. C) Averaging of fitted membrane intensity fraction *m* of repeated DC z-scans of cytoplasmic HIV-1 MA-EGFP. The uncertainty *s*_*m*_ in the fitted *m* value decreases as the square root of the number of repeated scans *n*.

To compare this simulation result with experiments, we performed DC z-scan measurements in cells co-expressing HIV-1 MA-EGFP and mCherry. Each z-scan was repeated 12 times at a single location within the cell to determine the mean and standard deviation of *m* from fits to each trace. We rescaled the experimental standard deviation to our reference conditions mentioned above 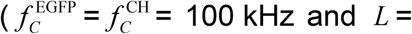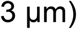 to simplify comparison. The rescaled experimental uncertainties (Fig. 5B) are generally in good agreement with the simulations. This validates the utility of our DC z-scan simulations and demonstrates that Eq. 15 provides a rudimentary, but reliable tool for gauging the expected uncertainty in recovering membrane populations from DC z-scan experiments.

These simulations suggest that a typical DC z-scan profile acquired in a cell with a cytoplasmic intensity of ~100 kHz can recover *m* within an uncertainty of ~1%. Because subsequent z-scan profiles acquired at the same location are independent statistical samples, we expect that averaging of multiple estimates of *m* reduces the uncertainty by 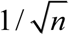, where *n* represents the number of scans. We collected repeated (*n* = 120) z-scans in cells co-expressing HIV-1 MA-EGFP and mCherry. These scans were grouped into sets of size *n*, followed by a calculation of the standard deviation of *m* over each set, which demonstrates the expected decrease in *s*_*m*_ as a function of averaging (Fig. 5C). Based on this result we chose to acquire 60 *s* (*n* = 12) of repeated z-scans and present the average and SEM estimated from these repeated passes to summarize the z-scan result for a given location in a cell.

The above results support the potential of DC z-scan as a method to detect weak interactions of proteins with the PM. We next tested the technique on cytoplasmic proteins without binding affinity to the PM. These measurements serve as controls to check the robustness of the method in the context of live cell measurements. For these data we only consider *m* from the top membrane and discard *m* from the bottom membrane for reasons that will be discussed later. As our first control, we performed repeated DC z-scans in the cytoplasm of U-2 OS cells expressing EYFP. The red-shift in the spectrum of EYFP as compared to EGFP increases the crosstalk (**α**^**YFP**^ =(0.80, 0.20)) into the red emission channel. The DC z-scan was fit to Eq. 13 with a crosstalk of zero, which is equivalent to treating the EYFP crosstalk signal as our cytoplasmic reference. The recovered membrane intensity fractions from many cells were histogrammed (Fig.6A) and describe a distribution mean of *m* = 0.08% ± 0.05% (SEM, *n* ~ 400), indicating that the DC z-scan procedure recovers a PM fraction that agrees with the expected result of 0% within 1.6 SD. In this control the fluorescent light in the reference channel originated from the same molecules as the fluorescence light detected by the green channel.

As an additional control we measured repeated DC z-scans in the cytoplasm of cells co-expressing EGFP and mCherry, which were fit to Eq. 13 with EGFP crosstalk (*α*^EGFP^ = 0.11). The histogram of the recovered membrane intensity fractions (Fig. 6D) has a mean of 0.32% ± 0.07% (SEM, *n* = 500), which implies a minute fraction of PM-bound EGFP and is in contradiction to the known properties of EGFP. We further examined a fusion construct linking EGFP and mCherry (EGFP-mCherry) as a third control (Fig. 6G). Repeating the same experiment resulted in a mean of –0.18% ± 0.03% (SEM, *n* = 350), which is 5.5 SD from the expected value of zero. These observations indicate the presence of small systematic errors that limit the determination of *m* values close to zero. We introduced a threshold of *m*_limit_ = 0.5% to account for the bias observed by the data of Fig. 6. Thus, we consider experimental *m* values that are within the range of ±*m*_limit_ to be consistent with *m* = 0 and only values that exceed *m*_limit_ count as a reliable indicator of PM binding. The recorded *m* values were graphed versus the cytoplasmic concentration of the proteins (Fig. 6) together with ±*m*_limit_. We further converted *m* and *m*_limit_ into the corresponding surface densities at the PM using Eq. 2 (Figs. 6C, F and I). The SEM of nearly all measured values overlaps with the range of ±*m*_limit_ for the negative control samples.

**Figure 6.**
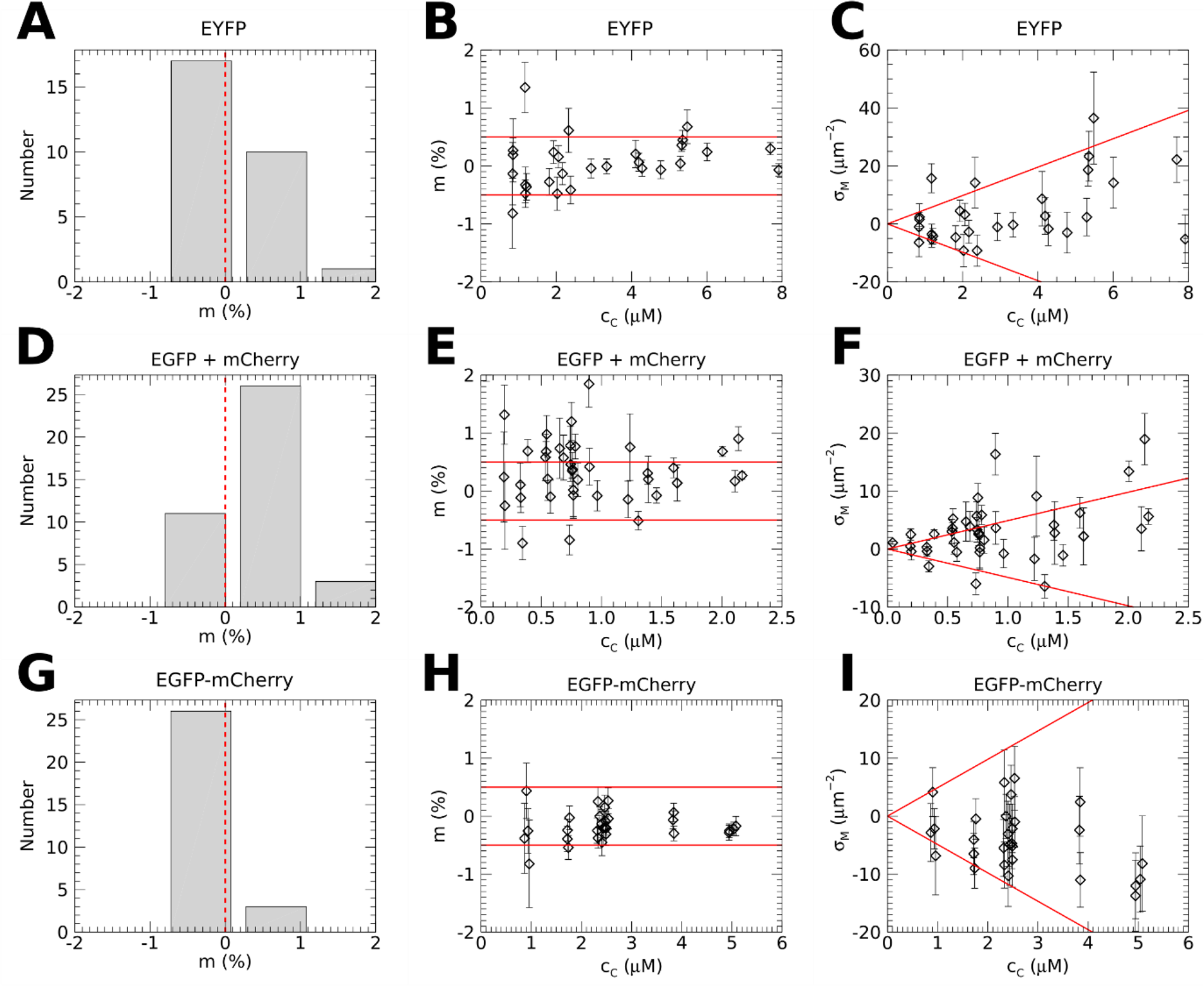
DC z-scan of controls with no PM binding. DC z-scans in cells expressing EYFP (top row), EGFP and mCherry (middle row) and EGFP-mCherry (bottom row). For each sample the PM binding from DC z-scan fits is presented as a histogram of the average *m* value from each z-scan location (A, D, G), as a scatter plot of average *m* vs cytoplasmic concentration *c*_*C*_ of the green channel (B, E, H), and as a plot of the surface density of PM binding *σ*_*M*_ vs *c*_*C*_ (C, F, I). The red solid lines denote ±*m*_limit_ and indicate the approximate region of null detection. Diamonds and error bars denote the average and SEM estimated from *n* = 12 repeated z-scans at each location within a cell.

During collection of the data we noted that DC z-scans frequently returned *m* values from the bottom PM that were significantly negative. Because the DC z-scan method introduced here is a relative measurement comparing an EGFP labeled protein against a soluble mCherry cytoplasmic marker, negative *m* values can arise if the red channel fluorescence trace contains a red emitting fluorescent contamination at the glass surface of the well plates (effectively adding a δ-layer at the bottom PM of the reference channel). We verified that this is the case for the well plates used in this study (Fig. S3) and thus we discarded *m* values associated with the bottom PM interface.

Although FRS2α is a major mediator of fibroblast growth factor signaling, its PM binding properties are not well understood. It has been reported that N-myristoylation of FRS2α is important for its localization to membranes and its ability to stimulate downstream signaling events (19). However, recent work by the Albanesi lab found that in addition to being myristoylated FRS2α is also double palmitoylated at C^4^ and C^5^, and that the palmitoyl groups are the main driver of PM binding (15). Because the FRS2α mutants employed in this study were at the detection limit of the SC z-scan method (14), we decided to revisit this study using DC z-scan to characterize their PM binding properties. WT FRS2α labeled with EGFP served as a reference to judge the influence of mutations on PM binding. Since FRS2α-EGFP strongly interacts with the PM, it was measured by SC z-scan in U-2 OS cells to determine its cytoplasmic and PM-bound concentration (*c*_*C*_ and *σ*_*M*_, respectively) as well as its membrane intensity fraction *m*. The PM binding is characterized by a plot of *m* vs *c*_*C*_ or alternatively by a plot of *σ*_*M*_ vs *c*_*C*_ (Fig. 7A & B). A global fit of both binding curves to a Langmuir-isotherm model (Eqs. 3 and 4) identified a dissociation coefficient *K*_d_ = 0.15 ± 0.05 μM and saturating surface concentration *σ*_0_ = 500 ± 80 μm^−2^ (Figs. 7A & B).

**Figure 7:**
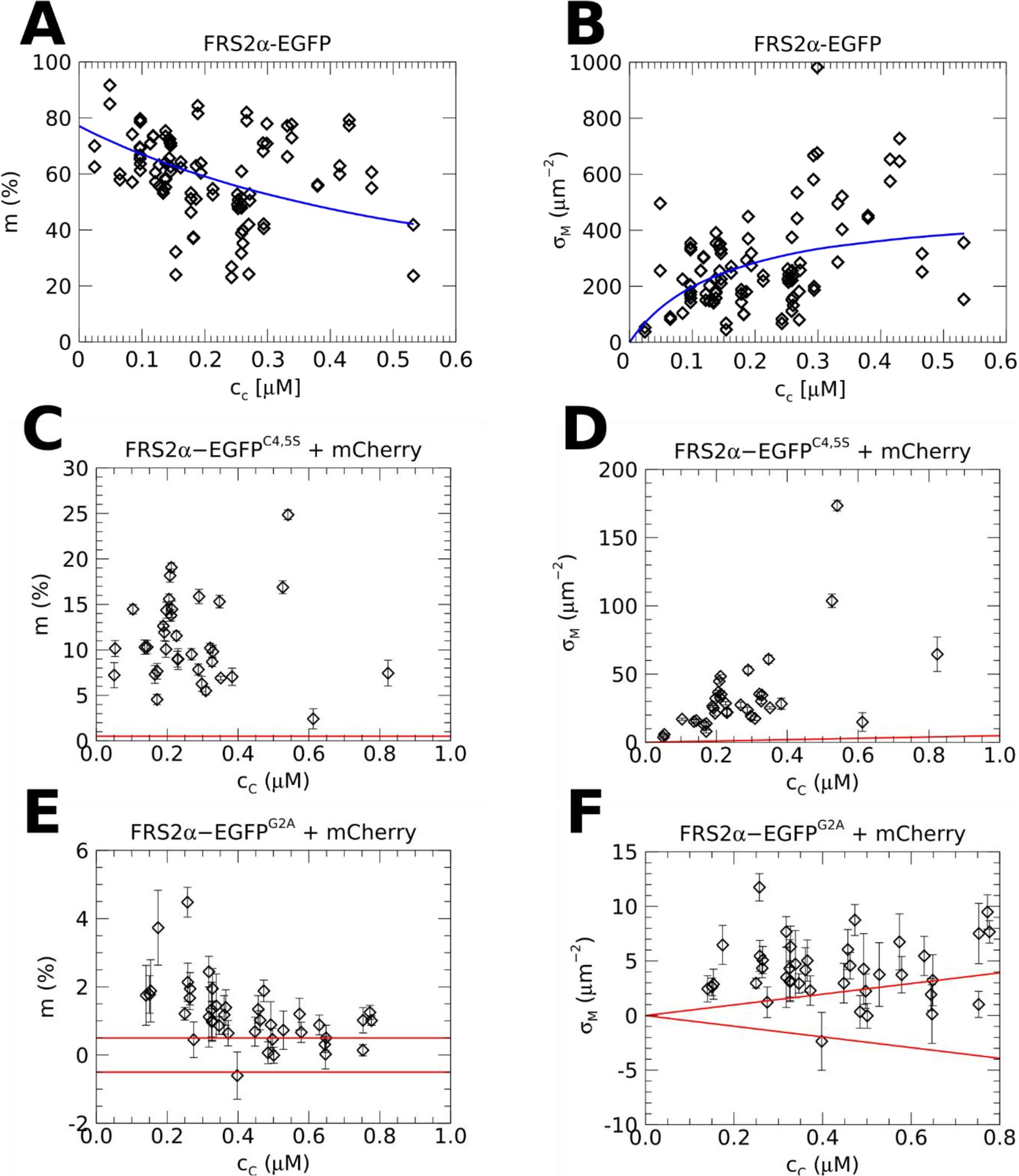
PM binding of WT FRS2α and two mutants. (A) *m* vs. *c*_*C*_ of WT FRS2α-EGFP as well as (B) *σ*_*M*_ vs. *c*_*C*_ together with fit to Langmuir-isotherm model (solid blue line). (C) *m* vs. *c*_*C*_ of FRS2α^C4,5S^-EGFP, and (D) *σ*_*M*_ vs. *c*_*C*_. (E) *m* vs. *c*_*C*_ of FRS2α^G2A^-EGFP, and (F) *σ*_*M*_ vs. *c*_*C*_. PM binding of FRS2α^G2A^-EGFP is detected below ~0.5μM and approaches the detection threshold of DC z-scan (red lines) at higher concentration.

Next, we performed DC z-scan measurements on the palmitoylation-deficient FRS2α^C4,5S^-EGFP construct to determine *m* (Fig. 7C). The mutation resulted in a severe reduction in the membrane intensity fraction from an average of 60% to ~10% (Figs. 7A & C). Similarly, the surface density of FRS2α^C4,5S^-EGFP at the membrane is significantly reduced (Fig. 7D). Because the cytoplasmic concentration expressed by the FRS2α-based constructs is low, the data are insufficient for a fit to a Langmuir-isotherm. However, we note that the binding data for FRS2α^C4,5S^-EGFP are consistent with *K*_*d*_ much higher than the concentration range we were able to access. The ratio *K*_*d*_/*σ*_0_ is determined by the limiting membrane intensity fraction *m*_0_ at low concentrations (Eq. 6), which predicts a *K*_*d*_/*σ*_0_ value of 5.3 μm^−1^. We further were able to estimate the dissociation coefficient of the mutant by assuming that the density *σ*_0_ of available binding sites at the PM is not affected by the mutation. In this case, we predict a dissociation coefficient of ~4.3 μM, which is an approximate 30-fold reduction in membrane binding affinity due to elimination of palmitoylation. This estimate demonstrates that the palmitoyl groups are the main determinant of PM binding. In addition, we also measured the EGFP-labeled G2A mutant FRS2α^G2A^, which eliminates both myristoylation and palmitoylation (15), and constructed plots of *m* vs *c*_*C*_ and *σ*_*M*_ vs *c*_*C*_ (Fig. 7E & F). PM binding of FRS2α-EGFP averaged 0.9 ± 0.1% (SEM, *n*=500), which is slightly above the detection limit of the DC z-scan method and suggests that FRS2α retains a small PM binding even in the absence of lipidation.

We next applied DC z-scan to characterize the binding of the matrix domain of HIV-1 Gag at the PM, which is driven by hydrophobic interactions due to N-terminal myristoylation at G^2^ as well as electrostatic interactions between acidic lipids and the highly basic region (HBR) of MA (20, 21). Although binding of MA to membranes at high concentration is readily detected by in-vitro studies (22–29), we were previously unable to detect binding of HIV-1 MA-EGFP to the PM of live cells with the SC z-scan technique (14), suggesting that its membrane fraction is generally below 10%. We performed DC z-scan measurements in the cytoplasm of cells co-expressing HIV-1 MA-EGFP and mCherry. The resulting plot of *m* versus *c*_*C*_ in Fig. 8A showed clear PM binding at all concentrations and a plot of *σ*_*M*_ versus *c*_*C*_ showed a clear increase in binding density for *c*_*C*_ between 2 and ~30 μM (Fig. 8B). Notably, *m* values averaged 6.2% within this concentration range and only ~10% of all DC z-scans yielded *m* values >10%. To confirm that the PM binding signal is attributable to the expected PM targeting motifs in MA, we performed DC z-scan measurements on the myristoylation deficient MA^G2A^ mutant with a truncated HBR (MA^G2A-ΔHBR^, Fig 8C & D). These measurements failed to detect PM binding, with an average *m* value of 0.23% ± 0.08% (SEM, *n* = 1000), which is below the DC z-scan detection threshold *m*_limit_ and comparable to the *m* value measured for the EGFP : mCherry negative control (Figs. 6E and F).

**Figure 8:**
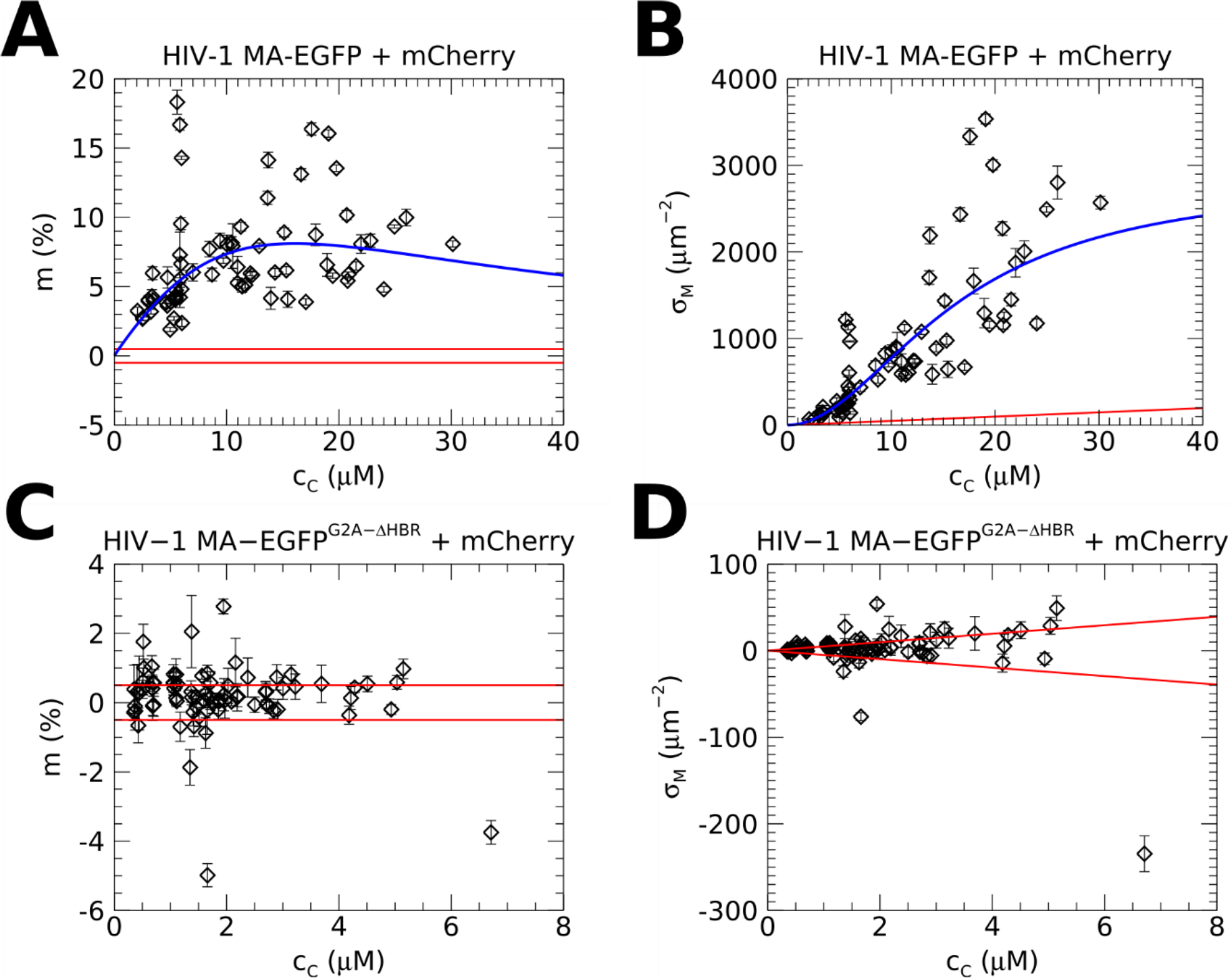
PM binding of HIV-1 MA and HIV-1 MA^G2A-ΔHBR^. (A) *m* vs. *c*_*C*_ for HIV-1 MA-EGFP. (B) *σ*_*M*_ vs. *c*_*C*_ for HIV-1 MA-EGFP. The solid blue lines represent a fit to the Hill equation (see Fig. S4). (C) *m* vs. *c*_*C*_ for HIV-1 MA^G2A-ΔHBR^–EGFP. (D) *σ*_*M*_ vs. *c_*C*_* for HIV-1 MA^G2A-ΔHBR^–EGFP. PM binding of HIV-1 MA^G2A-ΔHBR^–EGFP is generally below the detection threshold of DC z-scan (red lines).

Because a relatively wide concentration range was achieved for the expression of HIV-1 MA-EGFP, we further analyzed the DC z-scan results in Figs. 8A & B to investigate whether the PM binding can be described by a simple binding model. Notably, the *m* vs. *c*_*C*_ plot showed an increasing trend while the *σ*_*M*_ vs. *c*_*C*_ plot showed a convex curvature. Both of these shapes are incompatible with the Langmuir isotherm, which produces strictly decreasing and concave functions on the *m* vs. *c*_*C*_ and *σ*_*M*_ vs. *c*_*C*_ plots, respectively. We therefore fit the binding data from Figs. 8A & B to the Hill equation. As shown in Fig. S4, the Hill equation adequately described the measured HIV-1 MA binding curve with 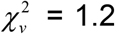 This fit recovered *K*_*d*_ = 16 ± 2 μM, *σ*_*0*_ = 2800 ± 400 μm^−2^ and Hill coefficient, *n* = 2.0 ± 0.08, indicating a significant degree of cooperativity in the PM binding of HIV-1 MA-EGFP. An additional fit to a Langmuir isotherm showed a poor quality of fit 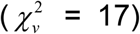, demonstrating that our qualitative observations that the PM binding of HIV-1 MA-EGFP cannot be described by a Langmuir isotherm model were recapitulated quantitatively.

## DISCUSSION

Our results demonstrate that DC z-scan significantly enhances the detection limit of protein binding to the PM. The key component of this improvement over SC z-scan is the presence of mCherry as a fluorescent paint to mark the cellular space, which provides an independent measure of the cytoplasmic thickness at the scan location. This additional information not only constrains the fit, which leads to a large reduction in the uncertainty *s*_*m*_ of the fitted membrane intensity fractions *m* (Fig. 5), but also removes the inherent bias of SC z-scan in recovering low membrane intensity fractions (*m* ≤ 10%, Fig. 4). Together, removal of bias and reduction in *s*_*m*_ improved the detection limit from ~10% for SC z-scan to below 1% for DC z-scan.

Quantitative z-scan analysis also relies on an optical system free of significant optical aberrations over the scan depth (30). In fact, we previously employed a co-expressed mCherry reference to interrogate PM binding of HIV-1 MA (31), but our earlier work was performed in the presence of chromatic and spherical aberrations. As a consequence, we were unable to rigorously analyze the DC z-scan traces in this earlier work and failed to detect HIV-1 MA PM binding. In this study we demonstrate that the use of two-photon co-excitation, the absence of pinholes in the detection path, and use of a high-quality water immersion objective permits a quantitative description of the DC z-scan traces sensitive resolution of PM binding. While the absence of statistical biases and chromatic effects in DC z-scan would in principle allow detection of even smaller membrane intensity fractions by reducing the uncertainty *s*_*m*_ based on averaging (Fig. 5C), we noticed the appearance of systematic errors at *m* ≤ 0.3% (Fig. 6). Although the source causing this effect is currently unknown, its amplitude is sufficiently small to render it negligible in most applications. We account for the potential presence of bias by setting a conservative detection threshold of *m*_limit_ = 0.5% for positive identification of PM binding. In practice we observed that slight fluorescent contamination at the coverslip surface is a far larger source for potential error in DC z-scan measurements. The fluorescence at the surface presents a signal not accounted for in the fit models, which biases the results. This is especially important at low expression levels, where the contaminating signal is strongest with respect to the sample. We expect that careful cleaning of the coverslip glass would eliminate or reduce the fluorescence at the interface, which should allow reliable DC z-scan data to be obtained from the bottom PM of adherent cells.

Utilizing DC z-scan enabled us to investigate the contributions of lipid anchors to PM binding of FRS2α. Because FRS2α is required for most aspects of fibroblast growth factor receptor (FGFR) signaling, and has itself been implicated in oncogenesis (32), mechanisms underlying its recruitment to the PM have been intensively investigated for over two decades. Previous measurements on the palmitoylation-deficient FRS2α^C4,5S^-EGFP construct was at the detection threshold of the SC z-scan technique and only was able to deduce the presence of residual PM binding by careful comparison with negative controls (15). In contrast, DC z-scan clearly identified a membrane intensity fraction of ~10% and was able to estimate a ~30 fold reduction of the binding affinity due to removal of the palmitoyl groups. FRS2α^G2A^-EGFP, which lacks all lipid anchors, showed a further reduction of PM binding. Unlike SC z-scan (15), DC z-scan was able to detect small amounts of residual PM binding of FRS2α^G2A^-EGFP, which may originate from constitutive binding between the FRS2α phosphotyrosine-binding domain and a juxtamembrane motif in endogenously FGFRs at the PM (33).

Extensive in vitro studies have reported binding of HIV-1 MA with various lipid substrates (22–28). The current work provides, to our knowledge, the first quantitative characterization of the PM binding of HIV-1 MA in-situ of the living cell. We found that MA binds to the PM of cells with a dissociation constant of ~16 μM, and saturating density of PM binding sites ~3000/μm^2^. Notably, the observed binding was cooperative in nature, which we speculate is a consequence of MA-MA oligomerization at the PM. Our observation of cooperativity is not unexpected. First, a recent SPR study measured a Hill coefficient of ~2 for binding to membranes composed of 100% DOPS lipids (26). Second, MA likely forms oligomers upon PM binding, which can dramatically increase PM affinity. This was previously shown by artificial dimerization of MA via addition of the HIV-1 capsid C-terminal domain, which increased the MA-membrane affinity by an order of magnitude (25). WT HIV-1 MA has been reported to trimerize variously as a 3D crystal (34), at high concentration in solution (35), when bound to biomimetic membranes (36) and in the mature viral particle (37). Trimerization of MA is thought to be important to accommodate the large cytoplasmic tail of the viral envelope protein for its proper packaging into infectious virus (37–39). Although the concentrations at which MA has been reported to trimerize in solution are significantly higher (~70 μM half point) than those probed by DC z-scan, PM binding likely favors oligomerization through increased MA-MA proximity due to partitioning into the reduced (effectively 2-dimensional) volume at the PM. Because of this membrane “scaffolding” effect, we expect trimerization of PM bound MA will occur at substantially lower cytoplasmic MA concentrations than are required to produce MA trimers in solution. We therefore suggest that after PM binding HIV-1 MA undergoes the expected trimerization reaction and that the trimeric form has significantly enhanced anchoring to the PM due to the increased binding energy associated with up to 3 MA-PM interfaces. This type of multi-step reaction is commonly associated with the observation of cooperativity. In such cases the Hill coefficient can be interpreted as a lower bound on the multiplicity of the oligomers (40, 41). We note that the Hill coefficient found for HIV-1 MA by DC z-scan is consistent with the expectation of MA trimers, although it does not exclude the possibility of a dimeric PM bound state, as we previously observed in the case of HTLV-1 MA (14).

The *K*_*d*_ ~16 μM measured in live cells by DC z-scan compares favorably to a recent NMR study that measured an apparent dissociation constant of 10 ± 2 μM for HIV-1 MA binding to phosphatidylinositol 4,5-bisphosphate (PIP_2_) containing liposomes (24). The SPR study of HIV-1 MA binding to tethered bilayers composed of DOPC/DOPS/PIP2/cholesterol yielded *K*_*d*_ values in the range of 0.8 to 2.4 μM depending on the specific lipid composition (26). However, the low salt conditions used in this study (50 mM; to prevent aggregation of purified MA) may have led to an exaggerated electrostatic component of MA-PM interaction and an increased MA-PM affinity. Increased PM affinity under low salt conditions has been reported in the case of Rouse sarcoma virus MA (42) and HIV-1 MA (25). Interestingly, the dissociation constant estimated by a previous SC z-scan study of HTLV-1 MA was ~200 nM (11), which differs from that of HIV-1 MA by nearly two orders of magnitude. These results serve to emphasize the striking diversity among retroviruses, particularly the orthoretroviruses, despite the relative conservation of retroviral MA domain structure.

Our study demonstrates that the DC z-scan approach eliminates statistical biases inherent to the SC z-scan method and furthermore increases the detection sensitivity by an order of magnitude. These advances significantly extend the range of cellular based PM binding assays. As demonstrated, DC z-scan allows evaluation of the contributions of individual PM binding motifs through mutation studies directly in the context of the living cell. In addition, DC z-scan successfully detected the weak binding of HIV-1 MA to the PM, and furthermore observed a cooperative binding process in the living cell. These results illustrate the potential of DC z-scan to contribute to our understanding of binding interactions between proteins and membranes in the authentic environment of the cell.

## AUTHOR CONTRIBUTIONS

I.A., S.R.K., L.M.M., J.P.A, and J.D.M were responsible for experimental design. I.A., S.R.K., J.H., and Y.C performed experiments, developed constructs and contributed analytical tools. I.A., S.R.K., J.H., and J.D.M. performed analysis. I.A., S.R.K., L.M.M., J.P.A, and J.D.M wrote the manuscript.

## ACKNOWLEDGMENTS

This work was supported by the National Institutes of Health grant, GM064589 (J.D.M, J.H., S.R.K., and Y.C.), and GM121536 (J.P.A., Y.C., J.D.M), and GM124279 (L.M.M., I.A., J.D.M.). I.A. was supported by NIH grant T32 AI083196.

